# Predicting transcriptional activation domain function using Graph Neural Networks

**DOI:** 10.1101/2024.05.08.593266

**Authors:** Farhanaz Farheen, Bradley K. Broyles, Yuanyuan Zhang, Nabil Ibtehaz, Alexandre M. Erkine, Daisuke Kihara

## Abstract

Analysis of factors that lead to the functionality of transcriptional activation domains remains a crucial and yet challenging task owing to the significant diversity in their sequences and their intrinsically disordered nature. Almost all existing methods that have aimed to predict activation domains have involved traditional machine learning approaches, such as logistic regression, that are unable to capture complex patterns in data or plain convolutional neural networks and have been limited in exploration of structural features. However, there is a tremendous potential in the inspection of the structural properties of activation domains, and an opportunity to investigate complex relationships between features of residues in the sequence. To address these, we have utilized the power of graph neural networks which can represent structural data in the form of nodes and edges, allowing nodes to exchange information among themselves. We have experimented with two kinds of graph formulations, one involving residues as nodes and the other assigning atoms to be the nodes. A logistic regression model was also developed to analyze feature importance. For all the models, several feature combinations were experimented with. The residue-level GNN model with amino acid type, residue position, acidic/basic/aromatic property and secondary structure feature combination gave the best performing model with accuracy, F1 score and AUROC of 97.9%, 71% and 97.1% respectively which outperformed other existing methods in the literature when applied on the dataset we used. Among the other structure-based features that were analyzed, the amphipathic property of helices also proved to be an important feature for classification. Logistic regression results showed that the most dominant feature that makes a sequence functional is the frequency of different types of amino acids in the sequence. Our results consistent have shown that functional sequences have more acidic and aromatic residues whereas basic residues are seen more in non-functional sequences.

## Introduction

The key factors for activation of eukaryotic genes are gene-specific activators. Each of these proteins contain two obligatory domains: DNA-binding domains and activation domains (ADs). DNA-binding domains provide gene specificity by interacting with specific DNA sequences, while ADs, within the same transcription factors, drive transcription initiation by orchestrating dynamic nuclear interactions. The DNA-binding domains have very specific conserved sequences which determine a variation of structure motifs, which in turn define the specificity of interaction with target promoter DNA sequence^1–3^. In contrast, ADs are highly variable in sequence, intrinsically disordered, and engage in fuzzy interactions with multiple often uncertain targets. The enigma of ADs stands for decades^4,5^. Recently with the advent of high throughput experimental approaches based on breakthroughs of massive parallel DNA synthesis and sequencing, AD analysis has been elevated on the new level.

The extremely high sequence variability of ADs, by some estimates >10^24^ sequence variants able to replace each other within the context of the same gene activator molecule^6–8^, make ADs an ideal target for machine learning (ML). Several attempts to develop ML models have been reported. The initial attempt based on the regression models allowed to define and test AD features which are important for the ML model performance^6^. Following the neural network (NN) based approach allowed to increase the accuracy of AD prediction^7^. However, the main reason for the higher accuracy of the NN model turns out to be the larger size of the dataset used for the ML training. When compared the regression model although slightly less accurate in prediction than NN model, allows better to see and develop ML features and to correlate them with the biochemical features of ADs^8^. Additional ML attempt using only sequences of natural transcription factors followed^9^, allowing to correlate ML with sequences existing in living cells. While the sequence features of ADs became clearer, understanding of structural AD features remains obscure. The recent availability of structure prediction methods such as AlphaFold^10^ and ESMFold^11^ as powerful tools for protein structure prediction allow their usage for the development of new ML models based on structural information, and the development and understanding of structural features of ADs. In this study we utilized ESMFold to convert the available dataset of annotated ADs^7^ into a dataset with ESMFold defined AD structures aiming to develop new ML models based on structural information and to gain information about structural features of ADs. ESMFold has aided in faster prediction of structures of the peptide sequences and with the large size of the dataset that we have used, this has helped in prompter analysis of results.

Although recently there have been attempts at predicting and analyzing activation domains using neural network architectures^7,9,12^, these have mostly involved Convolutional Neural Networks (CNN). While traditional CNNs are known to capture local patterns in the data, they may not always be able to handle long-range dependencies in sequences. Consequently, it becomes difficult to investigate into the connection or dependency between residues that may not be next to each other in the sequence. Moreover, while traditional ML methods such as logistic regression allow comprehension of the features that determine function of activation domains^8^, they may not capture the complex relationships between composition-based features or structure-based features.

In our work, we have aimed to utilize the advantage of Graph Neural Networks (GNN)^13^, which, in the last decade, has gained increasing popularity in the field of bioinformatics^14–18^. GNNs can represent data in the structure of a graph with nodes and edges, allowing them to harness a special ability called ‘message-passing’ that can permit adjacent nodes to share information among themselves. This is advantageous in the sense that it helps to capture information regarding dependency between nodes if they are connected by edges regardless of where they are positioned. In our study, we have used a dataset of more than 1 million peptide sequences to train and validate GNNs that are able to perform binary classification to determine whether a sequence is functional or not. For formulating the GNN, we have firstly followed the technique used in GNN-DOVE^15^, where the graph nodes represent atoms in the peptide structure. After this, we have developed a modified GNN containing a new graph formulation with residues as the nodes that allows residues even at large distances to exchange information among themselves. To identify the most prominent features that determine function in these peptides, we have also trained a logistic regression model. We have found that our GNNs are more accurate than plain logistic regression. Moreover, residue-level GNN outperforms atom-level GNN for classifying the peptides. We have experimented with several combinations of features and although secondary structure does not provide any meaningful contribution to the vanilla logistic regression model, addition of this feature in the residue-level GNN has led to the best performing model, compared to several other feature combinations that we experimented with, having accuracy, F1 score and AUROC scores of 97.9%, 71% and 97.1% respectively. We have shown that this model outperforms other existing neural network methods applied on this task^7,9,12^. Moreover, we have also analyzed whether an alpha helix being amphipathic has any contribution to AD function on a subset of the entire dataset and have observed that addition of this feature also improves the performance of the baseline residue-level GNN, indicating that the amphipathic property of a helix does have a meaningful contribution in determining functionality. Finally, through logistic regression, we have found that the most important feature that determines whether a sequence will be functional is the count or frequency of different kinds of amino acids in the sequence when compared to any other position-based or structure-based features.

## Materials and Methods

### Dataset

To train, validate and test our graph neural network and logistic regression methods, we used the Gcn4 dataset^7^ which has a total of 1,054,335 peptide sequences, each having a length of 30 amino acids. Among them, 37,923 sequences are labelled to be functional. We divided the dataset into train, validation and test samples with a 75:15:10 ratio to conduct our experiments. The train-validation-test split is shown in Table 1.

**Table 1:** Train-validation-test split of the dataset.

| Total samples | Training samples (75%) | Validation samples (15%) | Test samples (10%) |
| --- | --- | --- | --- |
| 1,054,335 | 790,751 | 158,150 | 105,434 |

### Graph Neural Network (Atom-level GNN)

To train the graph neural network, we first need to define the graph formulation. Since a graph is composed of nodes and edges, we will first define what these represent in our network. Let G(V,E) be a graph where V is the set of vertices or nodes and E is the set of edges of the graph. Two nodes are said to be adjacent if they are connected to each other by an edge. We can represent a graph’s edge using an adjacency matrix representation (A).

In the atom-level GNN network, each node represents each atom in the sequence. We have computed several features for these nodes, and they are listed in Table 2, the first five rows of which have been taken directly from the GNN-DOVE^15^ paper. The edges in the network represent the connection or bond between atoms as well as the distances between them. We have used two graphs following the method in GNN-DOVE which are denoted by G^1^ and G^2^ that have their edges represented by two adjacency matrices A^1^ and A^2^ respectively. In graph G^1^, only atoms that are connected by covalent bonds have edges between them to prioritize information from atoms that are connected and adjacent to one another in the 3D structure, whereas in G^2^, atoms within a short distance i.e. 10Å have edges connecting them to capture information from nearby atoms or locality. For G^2^, the atoms do not need to have covalent bonds between them to be connected by edges. This is done following GNN-DOVE architecture and in our case, it helps to prioritize information from atoms that are closer to each other. We can define the two adjacency matrices in the following way. Here *d_ij_* represents the distance between atoms i and j. The parameters μ and σ are learnable parameters with initial values set to 0 and 1 respectively.

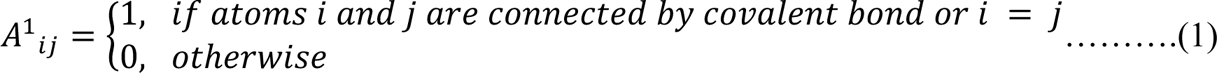

**Table 2:** List of features used in the Atom-level GNN. First five rows are taken from GNN-DOVE^15^ paper.

| Feature name | No. of columns for representation |
| --- | --- |
| Atom type (C,N,O,H,S,F,P,Cl,Br,B,H) | 10 |
| Degree or connections of atom (0,1,2,3,4,5) | 6 |
| No. of connected H atoms (0,1,2,3,4) | 5 |
| No. of implicit valence electrons (0,1,2,3,4,5) | 6 |
| Aromatic (0 or 1) | 1 |
| Position of residue the atom belongs to | 30 |
| Type of residue the atom belongs to | 20 |

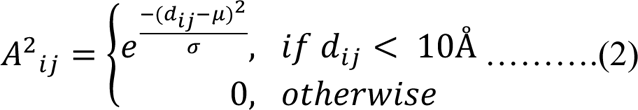

### Gate Augmented Mechanism with Attention

We then apply Attention and Gate-Augmented Mechanism in the same way as that of GNN-DOVE. Let us explain the Gate-augmented graph attention layer. If *x^in^* represents the node features, we can write it as: *x^in^ = {x_1_^in^, x_2_^in^, …, x_N_^in^}* where, *x* belongs to the real number space i.e. *x ε ℝ^F^*with *F* denoting the dimension of the node feature. At first, to retrieve the relative importance between the i-th and j-th node, the pure graph attention coefficient *e_ij_* is computed using the following set of equations:

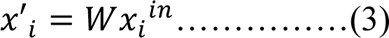

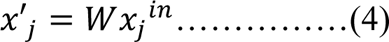

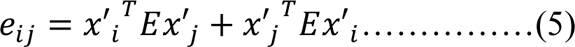

Equation 5 gives the pure graph attention coefficient. Here, W and E are learnable matrices. This coefficient is only computed for cases where we have positive values of *A_ij_*. To combine information from the pure graph attention coefficient with the adjacency matrix, we compute a normalized attention coefficient *a_ij_* in equation 6.

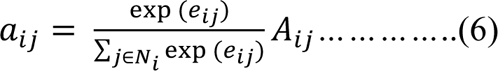

Here, *N_i_* represents the set of neighbors of i-th node. After this, we have calculated the updated node feature using the following equation:

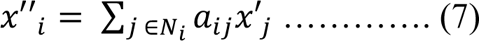

Equation 7 therefore, allows the consideration of node features. Finally, we have used the gate mechanism where we incorporate information from the input by inserting a residual connection. The gated graph attention is, thus, given in Equation 9.

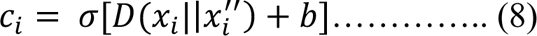

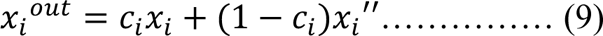

Equation 8 finds the coefficient value *c_i_* first. *σ* represents a sigmoid function. *D* and *b* are learnable parameters. The symbol || represents concatenation. Equation 9 gives the linear combination of *x_i_* and *x_i_’’.* If we denote this whole method involving attention and gate-augmented mechanism as gate-augmented graph attention layer (GAT), then we can use equation 10 to get the node embedding.

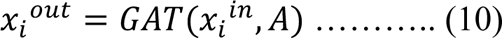

Since we have two adjacency matrices, and as we use a shared GAT for both of them, we will have two types of such node embeddings, *x^1^=GAT(x^in^,A^1^)* and *x^2^=GAT(x^in^,A^2^)*. To combine the information coming from adjacent and non-adjacent residues in the sequence, we added the embeddings of the two graphs. Note that equation 11 is different from the one in GNN-DOVE since in GNN-DOVE, *x^1^* was subtracted from *x^2^* to retrieve the information coming only from the intermolecular interactions. In our case, we want to combine the information from both adjacent and non-adjacent residues. So, the final output can be written as:

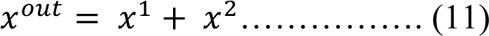

The GAT mechanism is done thrice iteratively i.e. the *x^out^* becomes *x^in^* and the whole process is repeated thrice after which we sum up the node embeddings for the whole graph to get the final representation which can be seen in equation 12.

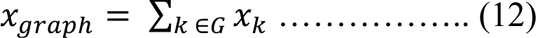

This *x_graph_* was then sent to a fully connected network with 4 layers and dimensions following the ones in GNN-DOVE^15^ (140 x 128 x 128 x 128). We have used RELU activation function between these layers. Finally, the output was sent through a sigmoid function to get one probability value between 0 and 1 which will represent the probability that a sequence is functional. Figure 1 shows the overall framework of atom-level GNN. The main idea has been borrowed from GNN-DOVE^15^. Two graphs are built from each peptide with atoms as nodes, and the final node embedding is obtained by adding the two node embeddings from the two types of graphs. This is done thrice iteratively after which the embedding is sent to a fully connected network. Figure 1 shows the dimensions of the layers. The final output is the probability value P which is between 0 and 1.

**Figure 1:**
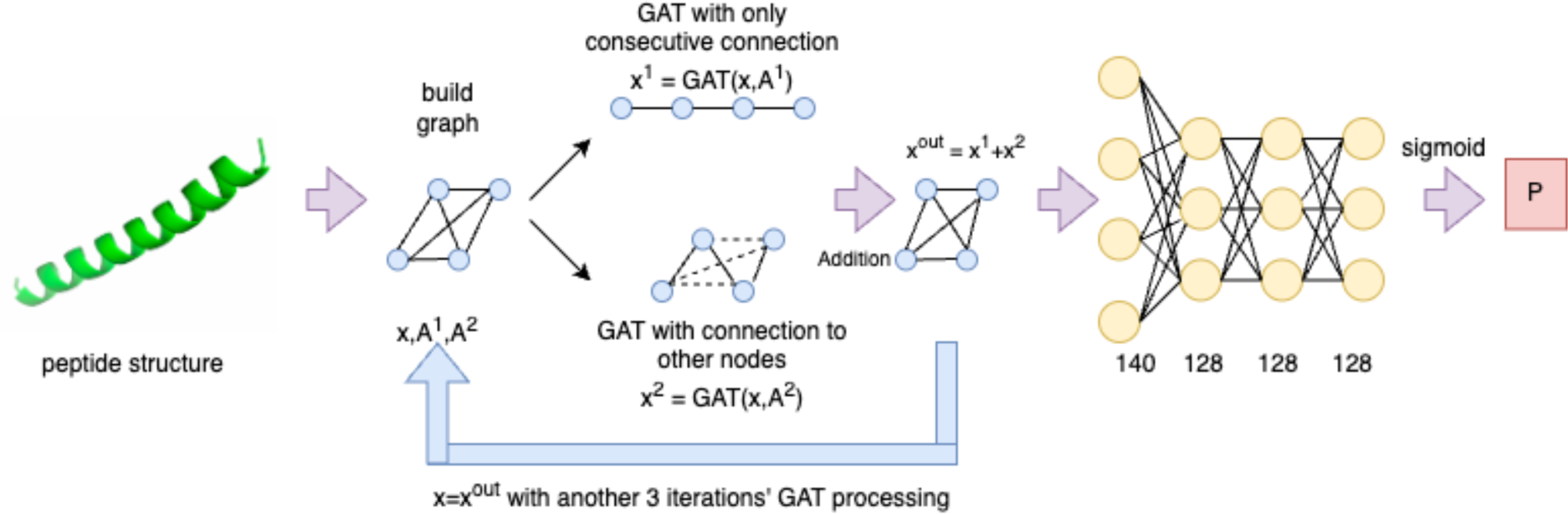
Framework of Residue-level GNN and Atom-level GNN. Two graphs are built from the peptide structure. After applying Gate Augmented Mechanism with Attention, node embeddings are added and sent to a fully connected network (FCN). Output from FCN is passed through a sigmoid function to get a final probability value P which is a value between 0 and 1.

### Atom-level GNN node features

The atom-level GNN has the following two differences when compared to the original GNN-DOVE: (1) After applying gate-augmented graph attention layer (GAT) on the input node feature *x_in_*, when we get two node embeddings, *x^1^=GAT(x^in^,A^1^) and x^2^=GAT(x^in^,A^2^*), they are added up to get the final embedding *x^out^* following equation 11, instead of subtraction in GNN-DOVE, since the goal in GNN-DOVE was to capture the information that only comes from the intermolecular interactions with other nodes in the protein complex model, whereas in our case, we are dealing with one peptide structure and want to combine the information from both adjacent and non-adjacent atoms; (2) Along with the GNN-DOVE features, we have also added two more types of features which consider the position of the atoms in the sequence and the type of residue they belong to. These features are listed in Table 2. The first five rows represent GNN-DOVE features and have directly been taken from the GNN-DOVE^15^ paper. They represent composition-based features of the atoms. The sixth feature is a new one and it represents the position of the residue that this current node or atom is in. Since there are 30 residues in each sequence, there can be 30 possible values and therefore this feature needs 30 columns for representation as it is one-hot encoded. The final feature represents the type of residue this atom belongs to. Since there are 20 different types of amino acids, there can be 20 possible values for this feature.

### Graph Neural Network (Residue-level GNN)

In the residue-level GNN network, the nodes represent the C-alpha atoms in the amino acids. Since all the sequences in our dataset have a length of 30^7^, there are 30 nodes, each representing an amino acid residue’s C-alpha atom. These nodes have features, which are listed in Table 3 and described in detail in the next subsection. On the other hand, the edges represent the connection and distances between the residues in the sequence. Following the technique used in GNN-DOVE^15^, we define two graphs G^1^ and G^2^ which have their edges represented by two adjacency matrices A^1^ and A^2^ respectively. G^1^ represents the graph where only adjacent residues are connected to each other. Two residues are adjacent if their indices in the sequence are consecutive, and this has no connection to the physical distance between them. Therefore, A^1^ has only binary values. On the other hand, graph G^2^ is a graph that takes into consideration the Euclidean distance between the three-dimensional coordinates of the residues. We have updated the adjacency matrix definition, and the changed equations are given in equation (13) and equation (14).

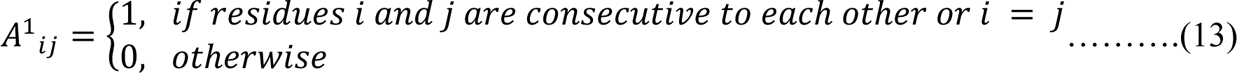

**Table 3:** List of features used in the Residue-level GNN.

| Feature name | No. of columns for representation |
| --- | --- |
| Position of residues | 30 |
| Type of amino acid | 20 |
| Type of residue (acidic, basic, aromatic) | 3 |
| Secondary structure (helix,beta,coil) | 3 |
| pLDDT | 7 |
| Relative accessible surface area | 10 |
| Amphipathic index | 6, 11, 24, 47 (depending on histogram bin-size) |

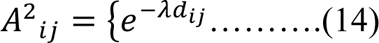

Here *d_ij_* is the distance between i-th and j-th residues. Here, *λ* is a learnable parameter and the initial value of *λ* is set to be 0. The idea of graph formulation has been borrowed from GNN-DOVE^15^ but this adjacency matrix *A^2^* has a different definition since in GNN-DOVE^15^, non-zero values were only considered when two residues were within 10 Å distance and the distance equation used there decayed for larger distances. In our case, we consider all 30 residues and change the decay formula to ensure that information from all other 29 residues is incorporated into the adjacency matrix, rather than only considering nearby residues. This is introduced in the residue-level GNN to capture information from all the residues instead of limiting the neighborhood within a certain locality.

After this, we apply the Attention and Gate Augmented Mechanism in the same way as we did in the atom-level GNN explained in the previous subsections. Figure 1 shows the overall framework of residue-level GNN. The main idea has been borrowed from GNN-DOVE^15^. Two graphs are built from each peptide with C-alpha atoms as nodes. The two node embeddings are added up and this process is repeated thrice before sending them to a fully connected network. The final output (P) is a value between 0 and 1.

### Node features (Residue-level GNN)

We have computed some node features, i.e. for a sequence, we apply some features to each residue. All features are one hot encoded. They are listed in Table 3 and described in detail below.

Since the length of the sequences is 30^7^, one residue can be assigned to one out of 30 positions and that’s what the first feature represents. It is one hot encoded which means we place 1 under the column that corresponds to that residue’s position. As there are 20 possible amino acids, the second feature has 20 columns and the residue will get 1 under its corresponding amino acid name, and the rest will be 0. For the third feature, we are only considering the appearance of some special residues in the sequence: acidic: Aspartic acid (D), Glutamic acid (E); aromatic: Tryptophan (W), Phenylalanine (F), Tyrosine (Y); and basic: Arginine (R), Lysine (K), Histidine (H). Since there are three such sub-groups (acidic, basic and aromatic), we have used 3 columns for this feature.

For secondary structure determination, we have used DSSP^19^ which gives 9 different types of structures as output. We have mapped these 9 to 3 being just alpha helix, beta sheet and coil, resulting in 3 columns for this feature representation.

We have used a structure prediction method (ESMFold^11^) for determining the structures of these sequences. This method also provides a confidence value which is a per-residue estimate of how confidently it predicted the structure. This is called the predicted local distance difference test (pLDDT). These values range from 0 to 100 where 0 means least confident and 100 means most confident. We have divided this range up into bins of size 10 giving 10 different bins (0-10,10-20,20-30….and so on). Since the dataset contained negligible samples with pLDDT less than 40, we have considered all values less than 40 to be one bin, which reduced the number of bins to 7 finally. For the relative accessible surface area feature, we have used DSSP’s output again which, along with secondary structure information, also outputs the accessible surface area values of the residues in the sequence. The accessible surface area is the surface area of the residue that is exposed or accessible to a solvent. Since the sizes of the residues can vary greatly depending on the type of amino acid, that’s why they are normalized with the help of a maximal solvent accessibility for each residue, the values of which have been taken from Table 1 of Rost et al. (1994)^20^. So, we can get the definition of relative accessible surface area from equation 15.

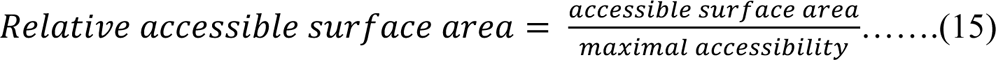

These values range from 0 to 100 where 0 means least exposed i.e. the residue is completely buried whereas, 100 means greatly exposed. We again divided this range up into bins of size 10, which gives us 10 bins in total and hence, this feature has 10 columns.

For the final feature, we have investigated the amphipathic index (AI) property of the data samples that have helices. An amphipathic alpha helix is basically a helix which contains both hydrophobic and hydrophilic residues arranged in a periodic manner such that one side of the helix becomes completely hydrophobic and the other becomes hydrophilic. The word ‘side’ here refers to two different faces of the helix that can be created if we look at the helical structure’s top view along its axis. We take the definition of amphipathic index of a helix from Cornette et al. (1987)^21^ and follow the method introduced in this paper to compute the amphipathic index of helices. According to Cornette et al. (1987), there are approximately 3.7 residues in one turn of an alpha helix, and therefore, we should notice a periodic variation in the hydrophobicity values of residues. This period should approximately be 3.7 residues per cycle. To detect this periodic variation, they calculate Fourier transform power spectrum using equation (16).

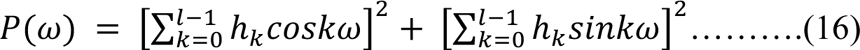

Here, *l* represents the length of the peptide sequence, *h_k_* denotes the hydrophobicity value for the k-th residue in the sequence and *ω* is the angular frequency. The hydrophobicity values have been taken from the PRIFT^21^ scale. Since hydrophobicity values are periodic in amphipathic helices, there should be a noticeable peak in the power spectrum. The amphipathic index is defined as the measure of how much of the power spectrum is concentrated around this peak. Therefore, the formula for AI can be written in equation (17).

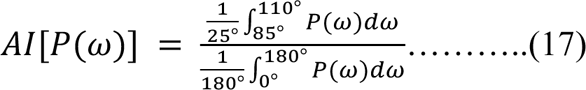

The width of the interval in the numerator is set to represent approximately the distance between half the maxima on each side of the peak of the spectra. Therefore, the AI value corresponds to how much of the area is centered about the expected peak of the spectrum compared to the total area under the spectrum. If a helix is amphipathic, this value is expected to be high, whereas, if it is not amphipathic, the numerator should be smaller and therefore the AI value should also be less. Since amphipathic index is a property of helix, we have at first considered sequences that have 10 consecutive residues that have been classified by DSSP to have a helical structure. There may be more than one such 10 length windows in a sequence. However, if there is no such window of 10 consecutive alpha helices in a sample, it has been removed from the dataset, resulting in a reduced dataset for particularly this feature since amphipathic index is specific to samples that have at least some forms of alpha helix. Note that this sort of formulation may cause one residue to belong to multiple windows of helix. Therefore, for each residue, we have taken the average AI of all the AI values corresponding to the different helix windows that the residue belongs to. So if we have a sequence *s = {s_1_,s_2_….s_30_}*, and if residues *s_1_* to *s_11_* are all part of a helix, then *s_2_* must be a part of two 10-length windows, the first being the window range (*w_1_*) of *s_1_* to *s_10_*, and the second (*w_2_*) being *s_2_* to *s_11_*. Then if we get two AI values *AI_1_* and *AI_2_* for *w_1_* and *w_2_* respectively, we will take an average AI value, 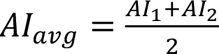. This *AI_avg_* will be the amphipathic index value corresponding to residue *s_2_*. Cornette et al. (1987) has mentioned 2 to be a reasonable cutoff meaning that it is more likely for a helix to be amphipathic if its AI value is greater than 2. We have then divided up the range of amphipathic index values to bins. Depending on the bin-size, the number of features may differ as well. For example, if the bin size is 1, then we only take the following bins: 0 to 1, 1 to 2, 2 to 3, 3 to 4, 4 to 5 and none (if the residue falls in no helix) which give us 6 columns for this feature. In this way, we take bin sizes of 0.5, 0.2 and 0.1 as well resulting in 10, 23 and 45 columns respectively.

Table 3 shows the number of samples of interest for this amphipathic index feature i.e. samples with at least 1 window of 10 consecutive helices. We can observe that, more than 50% of the data contains no such helix, and as a result considering the whole dataset would mean that majority of the samples will have zeros under all the feature columns. Consequently, we only considered a reduced subset of samples as shown in Table 4. We have simply filtered out the sequences with no 10-length helix from the train, validation and test sets.

**Table 4:** Number of samples of interest (at least one 10-length helix window) for amphipathic index feature analysis.

|  | No. of samples in the full dataset | No. of samples of interest (at least one 10-length helix) |
| --- | --- | --- |
| **Train set** | 790,751 | 225,597 |
| **Validation set** | 158,150 | 45,119 |
| **Test set** | 105,434 | 30,080 |
| **Total** | 1,054,335 | 300,796 |

### Logistic Regression

To understand the importance of different kinds of features, we trained a logistic regression model with 13 different kinds of features. Logistic regression is a simple network, and the features were computed on a sequence level, rather than a residue level. For each sequence, we computed both sequence and structure-based features. Structure prediction was done with ESMFold and to predict secondary structure, we used DSSP. We computed the set of features listed in Table 5.

**Table 5:**
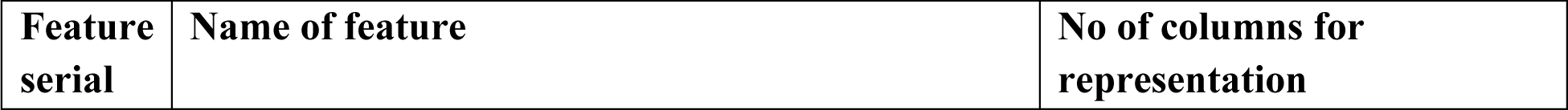
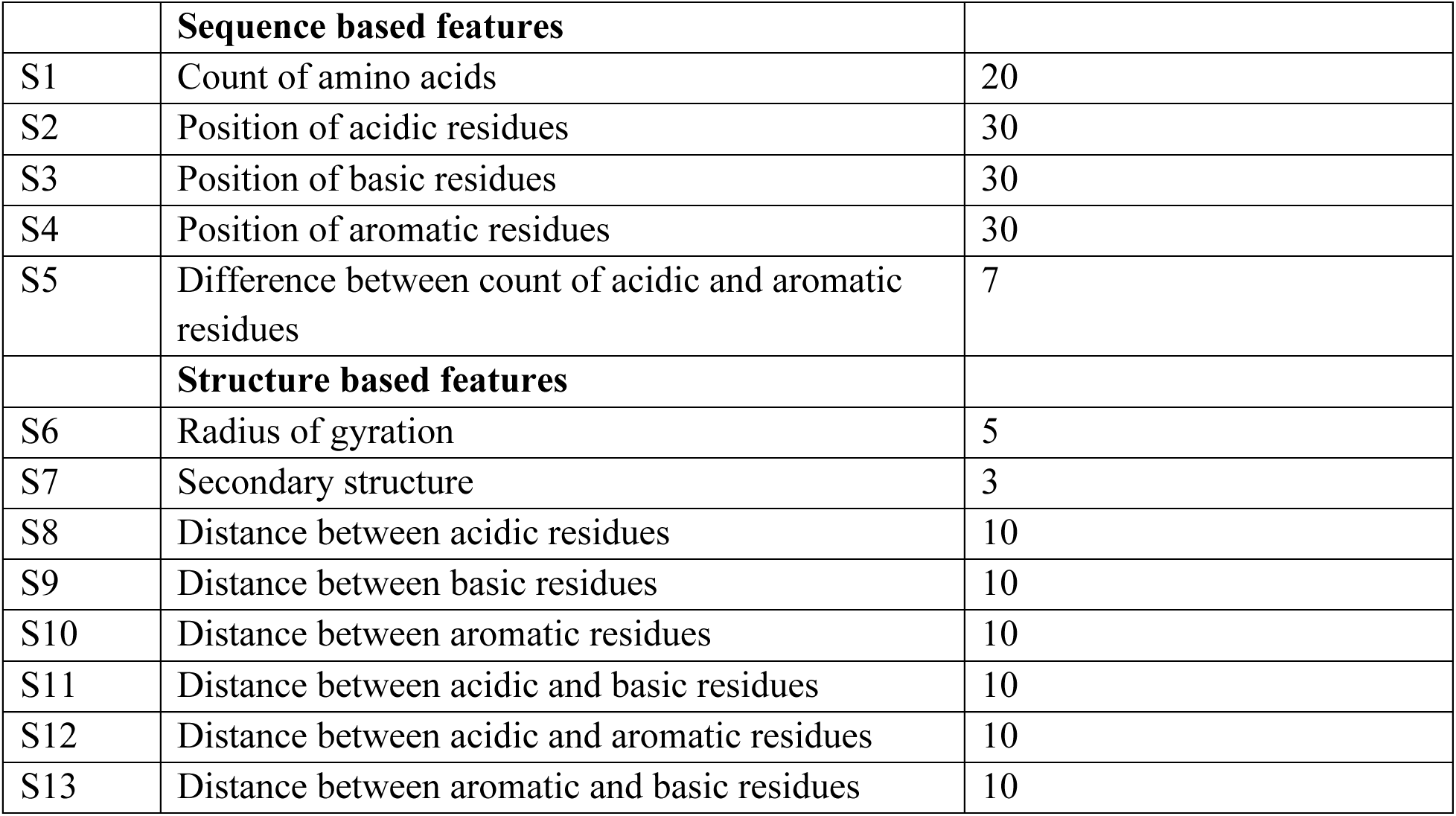
List of features used in the Logistic Regression model.

Logistic regression is a simple model and cannot handle complex representations of the data. We have computed features on a sequence-level for the regression model. There could be other features that are more suited for a residue-level or atom-level representation, but it will not be very meaningful to simply take an average of those embeddings to represent them on a sequence-level. Moreover, it was observed that the regression model achieved quite high accuracy (over 95%) with only sequence-based features and addition of more structure-based features was not improving the performance noticeably. Therefore, the model appears to have been saturated and adding more features may not contribute significantly.

There are 5 types of sequence-based features and 8 types of structure-based features. The first sequence-based feature is the count of amino acids feature. Since there are 20 different types of amino acids, this feature contains 20 columns, each corresponding to a different type of amino acid. We counted the number of each type of amino acid across the sequence and put that value under the corresponding column. For example, if a sequence has 3 Tryptophan (W), we put 3 under the column of W.

The second sequence-based feature represents the position of acidic residues. We have only considered the acidic residues for this feature which are Aspartic Acid (D) and Glutamic Acid (E). This feature contains 30 columns corresponding to 30 possible positions of the residues in the sequence. If a certain position contains any one of these 2 acidic residues, we assign 1 in that column, and 0 otherwise. The third and fourth features are computed in the same way as this one.

The difference is that we consider only basic residues in the third feature and aromatic residues in the fourth feature. By basic residues, we refer to Arginine (R), Lysine (K), and Histidine (H). By aromatic residues, we mean Phenylalanine (F), Tryptophan (W) and Tyrosine (Y). These two types of features were computed in the same way as that of the position of acidic residues and hence, they also contain 30 columns each.

The final sequence-based feature is the acidic-aromatic balance which refers to the difference in the count of acidic and aromatic residues in the sequence. We count the total number of aromatic residues (say *N_ar_*) and the total number of acidic residues (say *N_ac_*) in the sequence and find the difference i.e. *N_ar_ - N_ac_*. We divide this difference of count feature up into 7 bins or columns which are as follows: < -10, -10 to -5, -5 to < 0, 0, > 0 to +5, +5 to +10, > +10. The negative values mean that there are more acidic residues than aromatic residues and the positive values mean that there are more aromatic residues than acidic residues.

Now let’s come to the structure-based features. The first one is the radius of gyration feature. Radius of gyration is the root mean square distance of particles from axis. In our case, we consider the center of the peptide structure to be the axis. Let the center coordinate be *C(C_x_,C_y_,C_z_)* and let the coordinate of any atom be *P(P_x_,P_y_,P_z_)*. Then the distance between this atom and the center is the Euclidean distance between points *P* and *C* in the 3D coordinate system. The point *C* is obtained by taking the average coordinates of all the atoms in the peptide structure. Let *S* be the set of all atoms in one sequence and let the size of *S* be *N*.

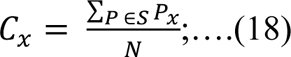

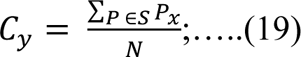

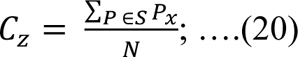

We can get the x, y, and z coordinates of C from equations 18, 19 and 20 respectively. The distance between a point P and C is simply the Euclidian distance between the two and can be written as *r(P,C):*

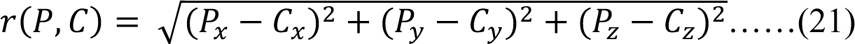

Next, we compute the Radius of gyration value for a sequence using equation 22.

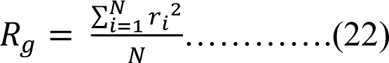

All these distance values are computed in Angstroms. The *R_g_* values are again divided up into 5 bins of length 5 each, which are: 5 to 10, 10 to 15, 15 to 20, 20 to 25 and 25 to 30 (there are no *R_g_* values greater than 30 or less than 5 for our dataset). A smaller *R_g_* value indicates that the protein is more compact, whereas a larger radius of gyration refers to a relatively longer peptide structure. The next 6 set of features are the distance-based features. We can divide them into two groups: the distance between same type of residues, and the distance between different types of residues. The first group includes the distance between acidic residues, distance between basic residues and distance between aromatic residues. For computing the distance between acidic residues, we first considered the coordinates of the residues which we consider acidic (Aspartic acid and Glutamic acid). Then we compute all pairwise Euclidean distances between all possible pairs of acidic residues. For example, if there are 3 acidic residues (say a,b and c) in the sequence, then we compute all three possible distances *distance(a,b), distance(b,c)* and *distance(c,a).* Then we put these distance values in histograms. We consider 9 such intervals, each of size 10 (0-10,10-20,….80-90) since there are no distance values greater than 90 and also consider an additional flag feature which receives binary values and will get 1 if there are no acidic residues at all, and 0 otherwise. For distance between basic residues, we follow the same technique considering only basic amino acids (Arginine, Lysine and Histidine) and for distance between aromatic residues, we compute the features in the same way considering the aromatic amino acids (Tryptophan, Tyrosine and Phenylalanine). We now come to the discussion about the second group of distance-based features which represent the distances between different kinds of residues. The first is distance between acidic and basic residues. We again take the coordinates of acidic residues and basic residues in the sequence and consider the pairwise distance between all possible pairs. For example, if there are 3 acidic residues and 2 basic residues then we have 3 x 2 = 6 possible pairs and hence, 6 possible distances. We again put them into bins of length 10 (0-10, 10-20,…..80-90) and keep a flag feature if either of acidic or basic residues is absent.

### Choice of structure prediction method

In both our GNN and Logistic Regression methods, we needed to use structure-based features to train the models. But our input is only the sequence. For structure prediction from sequences, we have used ESMFold. We opted to use ESMFold over the more popular AlphaFold for mainly two reasons: (1) ESMFold is a lot faster than AlphaFold and given our dataset size (∼1 million)^7^, we found this option more reasonable (2) We ran ESMFold and AlphaFold structure prediction methods on another set of 240 sequences (169 functional and 71 non-functional) that are also artificially generated. We then ran DSSP on these structures to get their secondary structure information. We considered a sequence to fall under the helix category if it had at least one window of 4 consecutive helices in its structure. It was seen that ESMFold predicted a lot more helices among the functional sequences compared to non-functional ones and this difference was seen to be statistically significant, whereas, for AlphaFold, no statistically significant result was seen. Table 6 gives the number of helices predicted by AlphaFold and ESMFold on the 240 sequences.

**Table 6:** Number of helices predicted by AlphaFold and ESMFold on the dataset of 240 sequences.

|  | AlphaFold | ESMFold |
| --- | --- | --- |
| **Functional (169)** | 24 helices | 59 helices |
| **Non-functional (71)** | 10 helices | 15 helices |

We wanted to see if these figures are statistically significant. Basically, we wanted to see if a certain sequence being functional has some connection with a structure prediction method predicting it to be helical or not. For that, we chose to perform Pearson’s chi-squared test where the null hypothesis is that there is no statistical significance between the number of helices in functional and non-functional sequences. For AlphaFold, we got a p-value of 0.98 which is greater than 0.05 showing that the result is not statistically significant. However, with ESMFold predictions, we got a p-value of 0.03 which is less than 0.05 showing that the result is statistically significant. Therefore, since we saw noticeable differences in the structures of functional and non-functional sequences predicted by ESMFold, we chose to predict the structures of our ∼1 million peptide sequences using this method.

### Training

For training the data, we had to resample the training set since the dataset is very imbalanced. To balance the dataset, we repeated samples from the positive or functional pool of sequences, and eventually sampled the same number of positive and negative samples. However, we did not balance the validation set or the test set since the inference should be done on a dataset that represents the true unsampled distribution of the data and testing on a balanced dataset may overestimate the F1 score.

For training the logistic regression model, we used L2 regularization or Ridge regression. For training Graph Neural Networks, we used the Adam optimizer and Binary Cross Entropy loss. We used a learning rate of 0.0001 for all the models and a dropout rate of 0.3 to avoid overfitting. For the residue-level GNN, we used a batch-size of 16,384 considering that the training set size is quite large. On the other hand, for training the atom-level GNN, we used a batch-size of 2048 given the computational resources. We used NVIDIA TITAN X (Pascal) (12GB), NVIDIA RTX A5500 (24GB), and NVIDIA RTX A6000 (48GB) GPU for running the GNN-related experiments. We trained different models with various feature combinations, and they converged after approximately 350 epochs.

### Post-processing to deal with low precision

Since we did not balance the test data, although the training was done on a balanced dataset, we observed very low precision and thus low F1-scores when we determined the predictions on our test set using a default threshold of 0.5 to classify the prediction as positive or negative. In binary classification, we need to set a threshold (which is usually 0.5) such that, if the predicted probability is greater than this threshold, the sample is classified as positive (or functional, in our case) and if the probability is smaller than the predetermined threshold, the sample is classified as negative (or non-functional). Adjusting the threshold is a matter of trade-off between precision and recall since, if we increase the threshold, fewer samples are classified as positive, leading to more false negatives and hence, lower recall. But if we decrease the threshold, fewer samples are classified as negative, leading to lower precision. Since the problem we primarily faced was low precision that resulted from a very imbalanced test set containing only 3.5% functional sequences, we decided to experiment with higher thresholds. We used the validation set for this task. For every epoch, the trained model was tested on the validation set and predictions were determined using a set of 10 thresholds – 0.5, 0.55, 0.6, 0.65, 0.7, 0.75, 0.8, 0.85, 0.9, and 0.95. For example, for 350 epochs, we saved 350 models and chose the combination of the model and the threshold that resulted in the highest F1-score on the validation set. Later the predictions on the test set were determined based on this final model that we chose after the threshold adjustment task.

## Results

### Evaluation metric

We used the following metrics for evaluating the performance of our models on the test set. Note that, samples that are truly functional and are predicted to be (1) functional, are true positives (2) non-functional, are false negatives. If they are truly non-functional and predicted to be (1) non-functional, they are true negatives (2) functional, they are false positives.

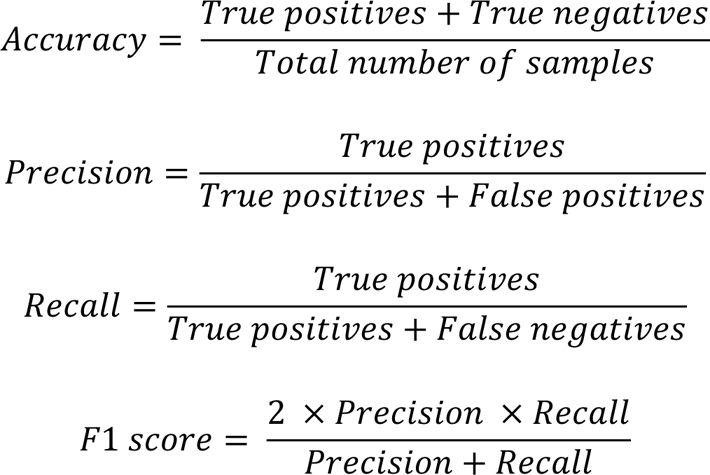

We have also used Area Under the Receiver Operating Characteristic (AUROC) curve as a performance metric. The Receiver Operating Characteristic curve is a graphical plot which is used to evaluate the performance of a binary classifier model. The X-axis denotes the False Positive Rate (FPR) and the Y-axis denotes the True Positive Rate (TPR). The FPR versus TPR values are plotted at different threshold points between 0 to 1. A curve whose area reaches 1 is a very accurate model that can differentiate between positives and negatives correctly at varied thresholds.

### Combining structure-based features with sequence-based features improves performance of logistic regression

For logistic regression, we have experimented with 3 kinds of models based on the type of features: (1) logistic regression with only sequence-based features, (2) logistic regression with only structure-based features, and (3) logistic regression with both sequence and structure-based features. The results are shown in Table 8 where it can be observed that only sequence-based features already perform very well, achieving 95.4% accuracy. In fact, if we compare only sequence-based features with only structure-based features, the former performs better for all metrics. This goes to show that the sequence alone has very powerful information that can distinguish between functional and non-functional samples. Compared to only structure-based features, they are a better classifier. However, if we combine all the sequence and structure-based features, the performance improves compared to the sequence-based features for all metrics. Although the accuracy improves very slightly – 95.4% to 95.5%, there is a decent increase in F1 score – 48.3% to 49.6% which goes to show that the combined model is better at handling the imbalanced test set. There is also a slight increase in the area under the ROC curve – 94.1% to 94.4%, telling us that the combined model can classify instances better at different thresholds. Both precision and recall are increased for the combined model, which means that both false negatives and false positives are reduced, leading to a better classifier overall when structure-based features are combined with sequence-based features. We can also view the confusion matrices in figure 2 to comprehend these results. Compared to figure 2(c), we can see more true positives and true negatives in figure 2(e). The combined logistic regression model classifies correctly 110 more samples that were not classified accurately by the sequence-only model. This goes to show that structural information is indeed relevant and useful for this classification task.

**Figure 2:**
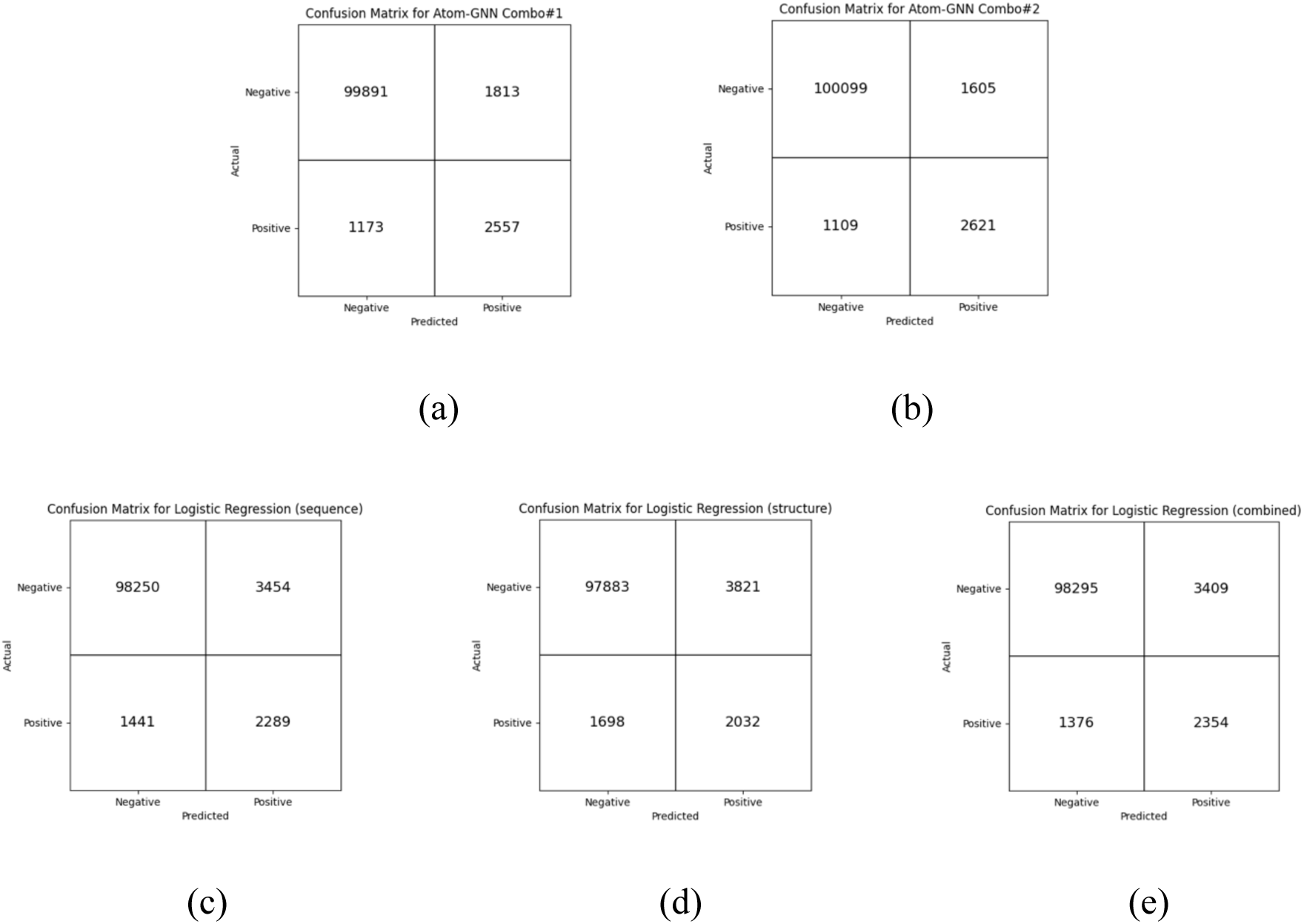
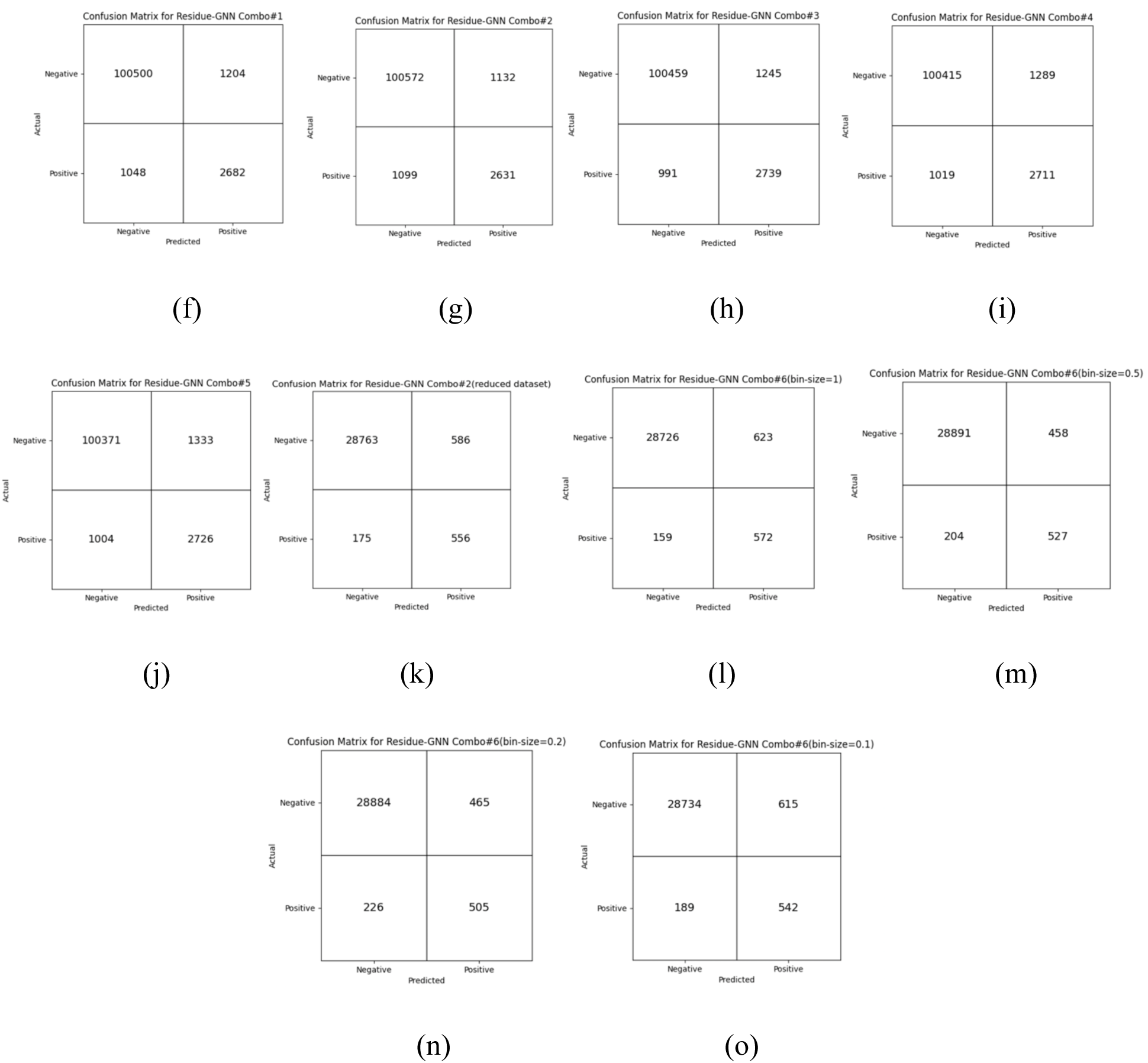
Confusion matrices for Atom-level GNN models (a,b), Logistic regression models (c-e), and Residue-level GNN models (f-o). The top left, top right, bottom left and bottom right cells in each matrix represent true negatives, false positives, false negatives and true positives respectively.

### Addition of secondary structure feature in residue-level GNN yields best results for all performance metrics

For experimenting with our residue level GNN model, we have trained models with different feature combinations as shown in Table 7.

**Table 7:**
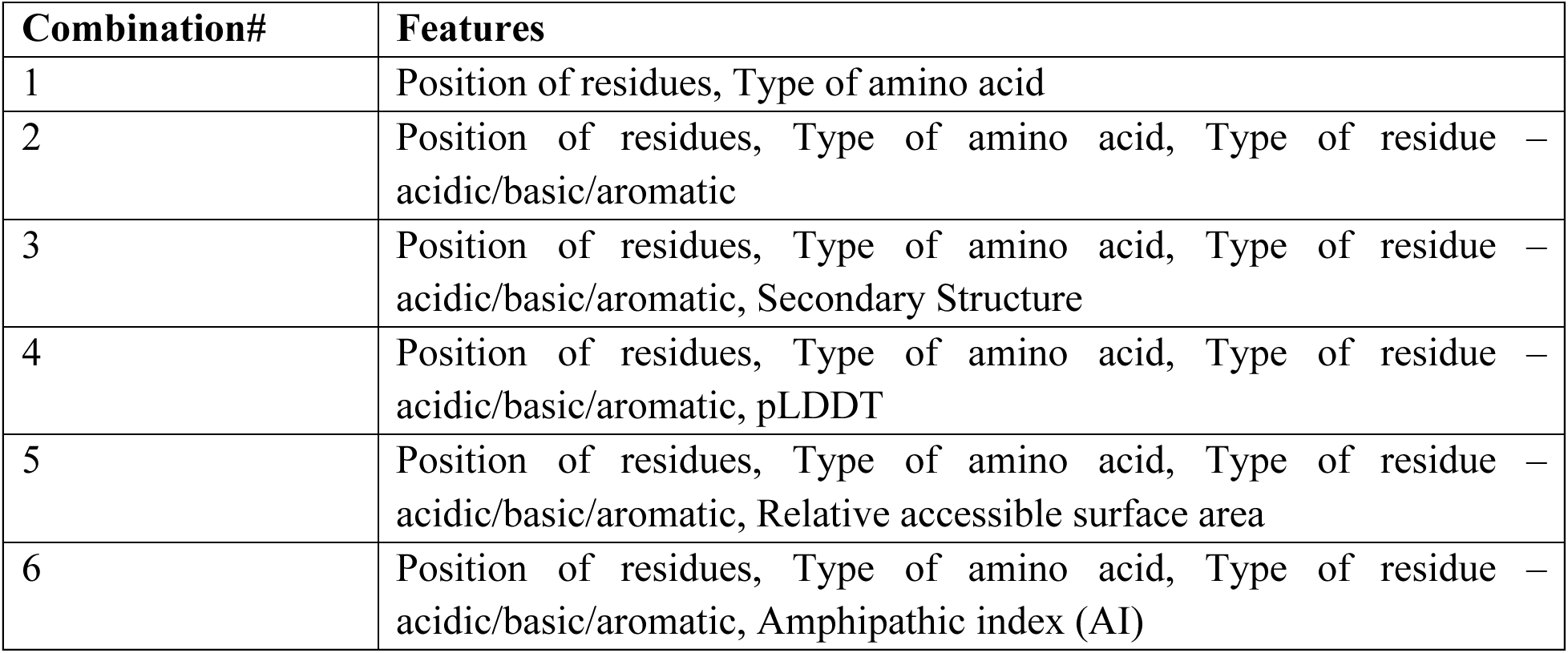
Feature combinations used in the experiments for determining the performance of residue-level GNN models.

The performance of these models (except row 6 which is shown in a different table – Table 10) are given in Table 8. We consider combination 2 with position of residues, type of amino acid, type of residue – acidic/basic/aromatic as the baseline and added other features to see if they improve performance any further. This combination already achieved really good performance, and although addition of pLDDT and accessible surface area did not improve the performance anymore, the addition of secondary structure as a feature improved the F1 score. Besides, we also experimented with the Amphipathic index feature on a subset of the dataset as explained in Table 4 of the methods section and compared it with our baseline model that was trained, validated and tested on the same set of samples. These results are shown in Table 10.

**Table 8:**
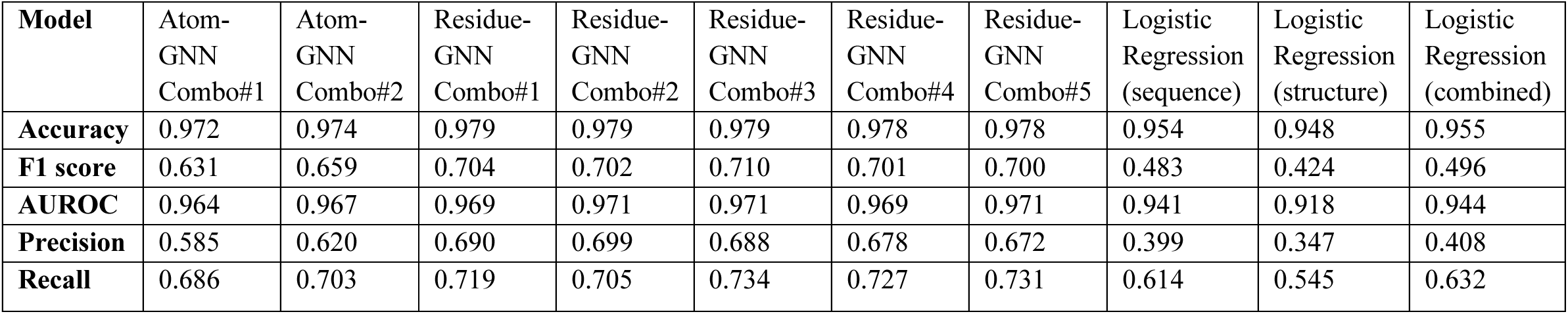
Performance of Atom-level GNN, Residue-level GNN and Logistic Regression models. Combination numbers are mapped to features in. **Tables 7 and 9**.

| **Model** | Atom-GNN<br>Combo#1 | Atom-GNN<br>Combo#2 | Residue-GNN<br>Combo#1 | Residue-GNN<br>Combo#2 | Residue-GNN<br>Combo#3 | Residue-GNN<br>Combo#4 | Residue-GNN<br>Combo#5 | Logistic<br>Regression<br>(sequence) | Logistic<br>Regression<br>(structure) | Logistic<br>Regression<br>(combined) |
| --- | --- | --- | --- | --- | --- | --- | --- | --- | --- | --- |
| **Accuracy** | 0.972 | 0.974 | 0.979 | 0.979 | 0.979 | 0.978 | 0.978 | 0.954 | 0.948 | 0.955 |
| **F1 score** | 0.631 | 0.659 | 0.704 | 0.702 | 0.710 | 0.701 | 0.700 | 0.483 | 0.424 | 0.496 |
| **AUROC** | 0.964 | 0.967 | 0.969 | 0.971 | 0.971 | 0.969 | 0.971 | 0.941 | 0.918 | 0.944 |
| **Precision** | 0.585 | 0.620 | 0.690 | 0.699 | 0.688 | 0.678 | 0.672 | 0.399 | 0.347 | 0.408 |
| **Recall** | 0.686 | 0.703 | 0.719 | 0.705 | 0.734 | 0.727 | 0.731 | 0.614 | 0.545 | 0.632 |

From Table 8, we can observe that, compared to the baseline model (combination 2), combination 3 which simply adds the secondary structure feature gives an increased F1 score of 71%. Its accuracy and AUROC are the same as that of the baseline but if we consider all three metrics – accuracy, F1 score and AUROC score, then combination 3 gives the best results in the whole table. This tells us that addition of secondary structure results in a model better at handling the imbalanced dataset, given its higher F1 score. It gives the best classifier among all the models that we experimented with on this dataset of more than 1 million sequences. Therefore, secondary structure feature is an important feature that can distinguish between functional and non-functional samples very well as shown by the residue-level GNN model, giving us a very high accuracy of 97.9%, F1 score of 71% and AUROC of 97.1%. This result can also be understood better from figure 2. We can observe that among all the confusion matrices in figure 2, the highest number of true positives can be seen in figure 2(h) which corresponds to combination 3. This means that this feature combination can identify the highest number of functional sequences in the dataset compared to any other model. Other features like pLDDT and accessible surface area have not improved the performance any further as shown in columns for combinations 4 and 5. Therefore, they have not been able to add any meaningful information to the binary classifier that we are developing.

### Both residue-level and atom-level GNN outperform logistic regression

From Table 8, we can see that, all residue-level GNN models outperform the logistic regression models. This is quite intuitive since logistic regression is a very simple model and it cannot capture the complex relationships between nodes – something that we can do using graph neural networks. GNN can aggregate information from neighboring or adjacent nodes and can capture non-linear relationships in the data as opposed to logistic regression. Although logistic regression already achieves a very high accuracy on this dataset – 95.5%, residue-level GNN can make it even more accurate with 97.9% accuracy. One very crucial thing to note is that logistic regression gives very poor F1 scores compared to residue-level GNN. In fact, the F1 scores are below 50%. This implies that the logistic regression models cannot achieve a good balance between precision and recall, and we have observed that they give very low precision. We can thus conclude that a simple model like logistic regression may appear to be very accurate but given a low F1 score on the imbalanced test set, it is obvious that they are not doing a good job in identifying the correct number of functional instances and are seen to generate many false positives. Besides, the accuracy and AUROC scores are also lower compared to residue-level GNN implying that the graph neural networks are superior for this classification task compared to logistic regression. This can also be seen in figure 2 where both the true positives and true negatives in figure 2(c-e) are fewer compared to the atom-level GNN models as shown in figures 2(a,b) and residue-level GNN models in figure 2(f-j) when trained and evaluated on the same dataset.

### Position of residues is an important feature as shown by the atom-level GNN results

For experimenting with the atom-level GNN, we tried two combinations of features which are shown in Table 9. The GNN-DOVE features correspond to the first five rows of Table 2.

**Table 9:** Feature combinations used in the experiments for determining the performance of atom-level GNN models.

| Combination# | Features |
| --- | --- |
| 1 | GNN-DOVE features, Type of amino acid |
| 2 | GNN-DOVE features, Type of amino acid, Position of residues |

Both atom-level GNN models give high accuracies, but atom-level GNN combination 2 outperforms atom-level GNN combination 1 for all performance metrics as we can see from Table 8. Combination 2 simply adds the feature of position of residues to combination 1. Addition of this feature improves the performance of the GNN which means that out of the 30 possible positions, where a particular residue is situated is important in determining the functionality of these peptide sequences. We can also comprehend this result from the confusion matrices in figures 2(a) and 2(b). We can see more true positives and true negatives in figure 2(b) compared to figure 2(c). In fact, addition of the position feature allows the GNN to classify 272 more samples accurately, indicating that this is an important feature in determining functionality.

### Residue-level GNN outperforms atom-level GNN

Although both Graph Neural Network models show better performance compared to logistic regression, among the two types of GNN, residue-level GNN is seen to perform better than atom-level GNN. In fact, all the five residue-level models with different combinations of features perform better than both the atom-level models for all performance metrics as shown in Table 8. The observation implies that for this particular task, having residue-level nodes in the graph formulation is more suitable and provides enough information to classify the sequences. Having atom-based formulation provides additional information that appears to be redundant for our classification task. Moreover, in the atom-level GNN, we do not consider any atoms beyond 10 Å whereas in the residue-level GNN, we take into consideration all the residues in the entire sequence. This shows that information from residues that are distant is also important in the GNN which is rather intuitive since the peptide sequences are relatively small (only 30 residues). This can be clearly seen in figure 2 since any true positive or any true negative value among the residue-level GNN models as shown in the confusion matrices of figure 2(f-j) are better than the true positive and true negative values respectively for the best atom-level GNN model as shown in figure 2(b).

### Amphipathic index is an important feature which improves performance compared to baseline model

Depending on the size of the histogram bins, we have experimented with 4 different models for analyzing the Amphipathic index (AI) feature. These are shown in Table 10.

**Table 10:** Performance of residue-level GNN with feature combination 6 for different bin-sizes on the smaller subset of the original dataset.

| **Model** | Residue-GNN<br>(Combo#2) | Residue-GNN<br>(Combo#6)<br>bin-size = 1 | Residue-GNN<br>(Combo#6)<br>bin-size = 0.5 | Residue-GNN<br>(Combo#6)<br>bin-size = 0.2 | Residue-GNN<br>(Combo#6)<br>bin-size = 0.1 |
| --- | --- | --- | --- | --- | --- |
| **Accuracy** | 0.975 | 0.974 | 0.978 | 0.977 | 0.973 |
| **F1-score** | 0.594 | 0.594 | 0.614 | 0.594 | 0.574 |
| **AUROC** | 0.969 | 0.967 | 0.969 | 0.966 | 0.966 |
| **Precision** | 0.487 | 0.479 | 0.535 | 0.521 | 0.468 |
| **Recall** | 0.761 | 0.782 | 0.721 | 0.691 | 0.741 |

It can be observed that compared to the baseline model shown in the first column of Table 10, addition of amphipathic index feature improves performance with a bin size of 0.5 and 0.2. But with a bin size of 0.5, we get higher accuracy, F1 score as well as similar AUROC scores. Therefore, for this subset of the dataset, the best model considering all three metrics of accuracy, F1 score and AUROC is given by the model in the third column with feature combination 6 and bin size of 0.5. The bin sizes were adjusted to see if we can get any feature representation of Amphipathic index that can give us a better classifier and this has been achieved on the reduced subset of the original dataset, leading to an accuracy of 97.8% compared to the baseline model’s performance of 97.5%. The F1 score is seen to have an even more noticeable increase – from 59.4% to 61.4% which implies that it does a much better job at handling class imbalance in the dataset. We can thus conclude that the amphipathic index feature is an important feature which has a meaningful contribution in determining whether a certain peptide sequence will be functional or not. From figure 2, we can observe that the confusion matrix in figure 2(m) representing the addition of amphipathic index feature with bin-size 0.5 gives the highest number of true negatives among figures 2(k-o) which represent models trained and evaluated for analyzing the amphipathic index feature.

### Amino acid count is the most important feature according to logistic regression

With logistic regression, we have conducted a feature importance test analysis in two different ways. The first is where we trained models with individual features for all 13 kinds of features. The results achieved are given in Table 11. The second is where we trained 13 models again, but this time, we left 1 feature out every time to check the fall of accuracy for removing that type of feature. This result is shown in Table 12.

**Table 11:** Performance of 13 logistic regression models trained with individual features (Method 1). Serial numbers are mapped to features in Table 5.

| Model | S1 | S2 | S3 | S4 | S5 | S6 | S7 | S8 | S9 | S10 | S11 | S12 | S13 |
| --- | --- | --- | --- | --- | --- | --- | --- | --- | --- | --- | --- | --- | --- |
| Accuracy | 0.951 | 0.904 | 0.899 | 0.856 | 0.626 | 0.382 | 0.616 | 0.688 | 0.900 | 0.616 | 0.876 | 0.944 | 0.928 |
| F1 score | 0.450 | 0.234 | 0.219 | 0.143 | 0.093 | 0.079 | 0.091 | 0.102 | 0.218 | 0.093 | 0.072 | 0.354 | 0.103 |
| AUROC | 0.932 | 0.827 | 0.817 | 0.722 | 0.593 | 0.551 | 0.601 | 0.664 | 0.695 | 0.622 | 0.567 | 0.885 | 0.619 |
| Precision | 0.373 | 0.163 | 0.150 | 0.090 | 0.051 | 0.042 | 0.049 | 0.057 | 0.151 | 0.051 | 0.049 | 0.299 | 0.092 |
| Recall | 0.568 | 0.413 | 0.401 | 0.339 | 0.539 | 0.750 | 0.540 | 0.499 | 0.392 | 0.556 | 0.135 | 0.435 | 0.117 |

**Table 12:** Performance of 13 logistic regression models trained by leaving 1 feature out every time (Method 2). Serial numbers are mapped to features in Table 5.

| Model | No S1 | No S2 | No S3 | No S4 | No S5 | No S6 | No S7 | No S8 | No S9 | No S10 | No S11 | No S12 | No S13 |
| --- | --- | --- | --- | --- | --- | --- | --- | --- | --- | --- | --- | --- | --- |
| Accuracy | 0.949 | 0.954 | 0.954 | 0.954 | 0.954 | 0.955 | 0.954 | 0.955 | 0.954 | 0.954 | 0.954 | 0.955 | 0.955 |
| F1 score | 0.437 | 0.489 | 0.491 | 0.492 | 0.496 | 0.497 | 0.493 | 0.496 | 0.495 | 0.496 | 0.487 | 0.495 | 0.495 |
| AUROC | 0.924 | 0.942 | 0.944 | 0.944 | 0.944 | 0.944 | 0.943 | 0.944 | 0.944 | 0.944 | 0.943 | 0.944 | 0.944 |
| Precision | 0.358 | 0.402 | 0.404 | 0.405 | 0.408 | 0.409 | 0.405 | 0.408 | 0.407 | 0.408 | 0.401 | 0.408 | 0.409 |
| Recall | 0.561 | 0.624 | 0.625 | 0.626 | 0.632 | 0.633 | 0.631 | 0.633 | 0.633 | 0.632 | 0.621 | 0.630 | 0.629 |

From Table 11, we can see that the most important feature is the count of amino acids which already achieves an accuracy of 95.1%. The combination of all the 13 types of features gives us 95.5%. This goes to show that the amino acid count itself contributes to almost the entirety of the accuracy achieved. On the other hand, we can see that some features like the radius of gyration leads to a completely random model as it gives us only 38.2% accuracy. Another thing to note is that the F1 score for all models apart from the one with the count of amino acids feature is significantly low – not even 25%. Other features like difference between count of acidic and aromatic residues, secondary structure, distance between acidic residues and distance between basic residues do not give very good accuracy with all of them being in a range of 61-62%. Although addition of secondary structure feature was seen to show good results in residue-level GNN, logistic regression could not capture the information from this feature well.

From Table 12, we can see one particular result which is consistent with Table 11. The fall of accuracy is the largest when we remove the count of amino acids feature. In order to evaluate the consistency between these two methods to judge the importance of features, we plotted a correlation graph of the two methods shown in figure 3. It shows us that both the methods agree on the amino acid count feature being the most important one. Although there are some other features that show decent accuracy by Method 1, they do not indicate high importance by Method 2 as the fall of accuracy is negligible when those features are removed. Clearly, the amino acid count feature dominates over the others in contributing to a high accuracy. Therefore, we have analyzed this feature in detail in the following subsection.

**Figure 3:**
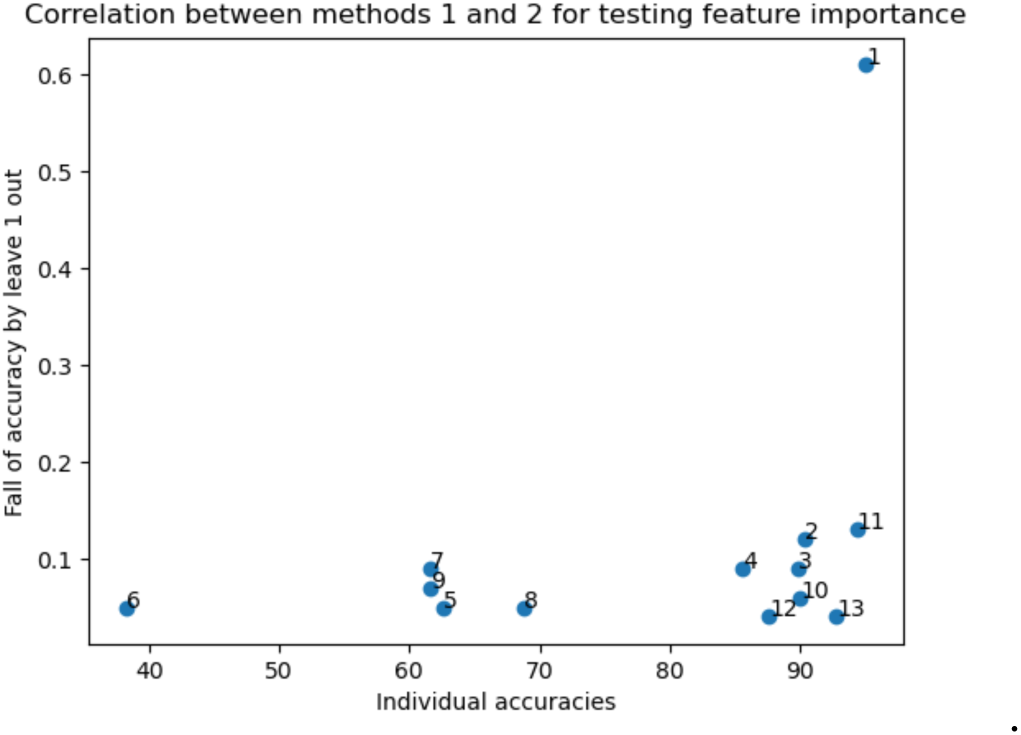
Correlation between two methods for feature importance test. X axis denotes the accuracies achieved by models trained with 13 types of features individually (Table 11). Y axis denotes the fall of accuracy when each of the 13 features are removed, one at a time (Difference between combined accuracy – 95.5% and accuracy values in Table 12). Serial numbers are mapped to features in Table 5. Data point 1 at the top right corresponds to the amino acid count feature.

### Aromatic (W,F,Y) and acidic (D,E) residues are important for functionality whereas basic (R,K,H) residues have the opposite effect

Since the amino acid count feature is predominantly the most important and crucial feature as shown by both methods 1 and 2 conducted for testing feature importance by logistic regression, we have analyzed this feature in more detail. The benefit of using logistic regression is that the feature coefficients obtained after training the model can allow us to determine which features contribute positively and which ones contribute negatively to the classification task. In logistic regression, the outcome depends on the sum of the products of features and their learned parameters or coefficients. The magnitudes of these coefficients determine how strongly the features contribute to the model and the sign of the coefficients determine whether they contribute positively or negatively. In our context, a positive sign means that the feature is usually noticed in functional sequences whereas a negative sign means that the feature is detrimental to functionality.

We have plotted the coefficients for the count of amino acids feature in figure 4 which are sorted based on their magnitudes. It can be observed that Tryptophan (W) has the highest positive impact on functionality whereas Arginine (R) has the most negative impact. Moreover, Lysine (K) is also seen to have a very large negative coefficient, indicating that R and K are usually seen in non-functional sequences. Another basic residue Histidine (H) also shows a negative coefficient but the magnitude is not as high compared to R and K. Acidic residues such as Aspartic acid (D) and Glutamic acid (E) as well as two other aromatic residues Phenylalanine (F) and Tyrosine (Y) have quite large positive coefficients which tell us that these residues are necessary for a sequence to be functional. All outcomes above are consistent with previously published results^6^.

**Figure 4:**
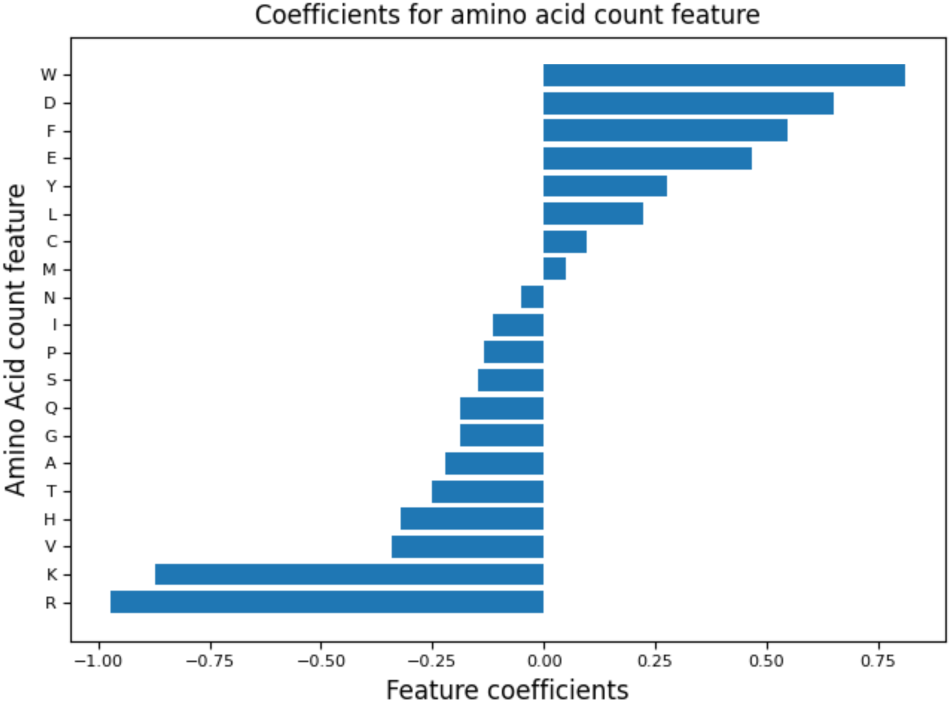
Coefficient values for the count of amino acids feature obtained by logistic regression. The X-axis here represents the coefficients whereas the Y-axis gives the amino acid 1 letter codes.

### Comparison with other methods

For comparing our model’s performance with other methods, we have tested three neural network architectures, all developed for predicting activation domains. The first is ADPred^7^ which was previously trained on the same set of sequences that we are using. They have provided as input the 30-length sequence to a convolutional neural network (CNN) followed by a dense neural network, which finally outputs a probability value. According to the suggestion of the authors, we have considered a threshold of 0.8 on the probability value of the 16^th^ residue to classify a sequence as positive or negative. After testing this method using our test set of 1,05,434 sequences, the accuracy achieved is 95.7% and the F1 score is 60.6% with an AUROC of 96.7%. Compared to this, our best model (residue-level GNN with feature combination#3) that gives accuracy, F1 score and AUROC of 97.9%, 71% and 97.1% respectively performs better. One thing to note is that their precision is quite low – 46.9% indicating that this method falsely predicts many non-functional sequences to be functional. The second method we compared with is PADDLE^9^ which also uses a deep convolutional neural network. Since PADDLE outputs a numerical value between -1 to 12, we have converted this into a classification task. According to the suggestion of the authors, we have considered both 4 and 6 as thresholds and to be stringent, we have also considered less than 4 to be negatives and greater than 6 to be positives. For all three cases, our method’s accuracy, F1 score and AUROC are better than those of PADDLE. These results are shown in Table 12. Another thing that we observe here is that the F1 score is less for PADDLE compared to both our method and ADPred.

**Table 12:** Comparison between our method and (1) ADPred and (2) PADDLE.

| **Model** | Residue-level<br>GNN<br>Combo#3 | ADPred | PADDLE<br>(threshold =<br>4) | PADDLE<br>(threshold =<br>6) | PADDLE<br>(neg < 4, pos<br>> 6) |
| --- | --- | --- | --- | --- | --- |
| **Accuracy** | 0.979 | 0.957 | 0.967 | 0.967 | 0.962 |
| **F1 score** | 0.710 | 0.606 | 0.337 | 0.140 | 0.163 |
| **AUROC** | 0.971 | 0.967 | 0.913 | 0.913 | 0.913 |
| **Precision** | 0.688 | 0.469 | 0.604 | 0.776 | 0.776 |
| **Recall** | 0.734 | 0.857 | 0.234 | 0.077 | 0.091 |

Besides, we have also compared our method with Mahatma et al. (2023)^12^ which also uses the dataset we have used, but a balanced one. Instead of experimenting with 1,054,335 sequences, they have taken all the 37,923 functional samples and appended 37,922 non-functional samples to create their dataset. They have also used CNN in their neural network architecture and combined it with two bidirectional long short-term memory (biLSTM) layers. They report an accuracy of 91.95% as well as F1 scores obtained using different architectures and parameters. The highest F1 score they reported is 91.95%. To conduct a fair comparison with their method, we have evaluated our best model (residue-level GNN with feature combination 3) on a subset of our original test set. Our original test set contains 3,730 functional samples. In the new test set, we have included all these 3,730 positive samples and appended equal number of negative samples to create a balanced test set. These negative samples were randomly selected, and the task was conducted for 10 iterations so that we can allow different negative samples to be considered while testing. Every iteration thus resulted in a different test set with fixed positive samples but different negative samples. Since this test set is balanced, we used a default threshold of 0.5 to classify the sequences. The comparison results are shown in Table 13.

**Table 13:** Comparison between our method and Mahatma et al. (2023).

| **Model** | Mahatma<br>et al | Residue<br>GNN<br>Iteration<br>1 | Residue<br>GNN<br>Iteration<br>2 | Residue<br>GNN<br>Iteration<br>3 | Residue<br>GNN<br>Iteration<br>4 | Residue<br>GNN<br>Iteration<br>5 | Residue<br>GNN<br>Iteration<br>6 | Residue<br>GNN<br>Iteration<br>7 | Residue<br>GNN<br>Iteration<br>8 | Residue<br>GNN<br>Iteration<br>9 | Residue<br>GNN<br>Iteration<br>10 |
| --- | --- | --- | --- | --- | --- | --- | --- | --- | --- | --- | --- |
| **Accuracy** | 91.95% | 92.71% | 92.84% | 92.96% | 93.22% | 93.15% | 93.06% | 93.10% | 92.86% | 93.14% | 92.95% |
| **F1 score** | 91.95% | 92.68% | 92.81% | 92.92% | 93.16% | 93.10% | 93.01% | 93.05% | 92.82% | 93.08% | 92.91% |

It can be observed from Table 13 that, for all iterations, our method outperformed Mahatma et al. (2023) in terms of both accuracy and F1 score. This attests to our method’s advantage over others even if we consider a balanced dataset for this classification task.

### Conclusion

We have utilized a Graph Neural Network for identifying sequences that are functional transcriptional activation domains and achieved highly accurate models. Analysis of different feature combinations has allowed us to judge which properties of these sequences and structures have meaningful contribution to AD function. Although there has been some investigation into the impact of secondary structure on the functionality of these peptides^7,9,12^, our method extensively analyses several structure-based features apart from only secondary structure as properties of individual residues and atoms. We have been able to achieve a performance better than other existing methods and have also identified the most important feature through a logistic regression model.

Our experiments have revealed that the secondary structure feature does have a meaningful contribution in classifying functional sequences as it helps to achieve a higher recall i.e. fewer false negatives and addition of this feature gives us the best performing model for all the performance metrics – accuracy, F1 score and AUROC. This goes to show that whether a peptide sequence is a functional transcriptional activation domain or not does depend on the secondary structures of the peptide i.e. whether the residues fall into a helix, beta or coil structure. Moreover, the results concerning amphipathic index suggest that the hydrophobic and hydrophilic nature being present on the opposite faces of the peptide also affects the function of the peptides. We have seen that the most important feature that distinguishes whether a peptide will be functional or not is the frequency of amino acids. After analyzing this feature in greater detail, we have found that the presence of acidic and aromatic residues is necessary for the peptide to be a functional transcriptional activation domain. On the other hand, presence of basic residues is detrimental to the function of the peptides. As future work, we can focus on analyzing some more structure-based features.

## Acknowledgement

This work was partly supported by funding from the National Science Foundation, MCB1925643. DK also acknowledges support from National Science Foundation (DBI2003635, DBI2146026, IIS2211598, DMS2151678, and CMMI1825941) and by the National Institutes of Health (R01GM133840).

## Code Availability

Our codes are available at https://github.com/kiharalab/GNN-TAD for free academic use.

## Notes

### Competing Interest Statement

The authors have declared no competing interest.

## References

1. Garvie, Colin W., and Cynthia Wolberger. “Recognition of specific DNA sequences.” Molecular cell 8.5 (2001): 937–946.

2. Luscombe, Nicholas M., et al. “An overview of the structures of protein-DNA complexes.” Genome biology 1 (2000): 1–37.

3. Hossain, Mir A., et al. “Artificial zinc finger DNA binding domains: versatile tools for genome engineering and modulation of gene expression.” Journal of cellular biochemistry 116.11 (2015): 2435–2444.

4. Strader, Lucia C., et al. “The complexity of transferring genetic information.” Molecular Cell 83.3 (2023): 320–323.

5. Erkine, Alexandre M. “‘Nonlinear’biochemistry of nucleosome detergents.” Trends in biochemical sciences 43.12 (2018): 951–959.

6. Ravarani, Charles NJ, et al. “High-throughput discovery of functional disordered regions: investigation of transactivation domains.” Molecular Systems Biology 14.5 (2018): e8190.

7. Erijman, Ariel, et al. “A high-throughput screen for transcription activation domains reveals their sequence features and permits prediction by deep learning.” Molecular cell 78.5 (2020): 890–902.

8. Broyles, Bradley K., et al. “Activation of gene expression by detergent-like protein domains.” Iscience 24.9 (2021).

9. Sanborn, Adrian L., et al. “Simple biochemical features underlie transcriptional activation domain diversity and dynamic, fuzzy binding to Mediator.” Elife 10 (2021): e68068.

10. Jumper, John, et al. “Highly accurate protein structure prediction with AlphaFold.” Nature 596.7873 (2021): 583–589.

11. Lin, Zeming, et al. “Evolutionary-scale prediction of atomic-level protein structure with a language model.” Science 379.6637 (2023): 1123–1130.

12. Mahatma, Saloni, et al. “Prediction and functional characterization of transcriptional activation domains.” 2023 57th Annual Conference on Information Sciences and Systems (CISS). IEEE, 2023.

13. Scarselli, Franco, et al. “The graph neural network model.” IEEE transactions on neural networks 20.1 (2008): 61–80.

14. Ioannidis, Vassilis N., Antonio G. Marques, and Georgios B. Giannakis. “Graph neural networks for predicting protein functions.” 2019 IEEE 8th International Workshop on Computational Advances in Multi-Sensor Adaptive Processing (CAMSAP). IEEE, 2019.

15. Wang, Xiao, Sean T. Flannery, and Daisuke Kihara. “Protein docking model evaluation by graph neural networks.” Frontiers in Molecular Biosciences 8 (2021): 647915.

16. Cao, Yue, and Yang Shen. “Energy-based graph convolutional networks for scoring protein docking models.” Proteins: Structure, Function, and Bioinformatics 88.8 (2020): 1091–1099.

17. Feng, Qingyuan, et al. “Padme: A deep learning-based framework for drug-target interaction prediction.” arXiv preprint arXiv:1807.09741 (2018).

18. Cai, Ruichu, et al. “Dual-dropout graph convolutional network for predicting synthetic lethality in human cancers.” Bioinformatics 36.16 (2020): 4458–4465.

19. Kabsch, Wolfgang, and Christian Sander. “Dictionary of protein secondary structure: pattern recognition of hydrogen-bonded and geometrical features.” Biopolymers: Original Research on Biomolecules 22.12 (1983): 2577–2637.

20. Rost, Burkhard, and Chris Sander. “Conservation and prediction of solvent accessibility in protein families.” Proteins: Structure, Function, and Bioinformatics 20.3 (1994): 216–226.

21. Cornette, James L., et al. “Hydrophobicity scales and computational techniques for detecting amphipathic structures in proteins.” Journal of molecular biology 195.3 (1987): 659–685.

